# The heparin-binding proteome in normal pancreas and murine experimental acute pancreatitis

**DOI:** 10.1101/497271

**Authors:** Quentin M. Nunes, Dunhao Su, Philip J. Brownridge, Deborah M. Simpson, Changye Sun, Yong Li, Thao P. Bui, Xiaoying Zhang, Wei Huang, Daniel J. Rigden, Robert J. Beynon, Robert Sutton, David G. Fernig

## Abstract

Acute pancreatitis (AP) is acute inflammation of the pancreas, mainly caused by gallstones and alcohol, driven by changes in communication between cells. Heparin-binding proteins (HBPs) play a central role in health and diseases. Therefore, we used heparin affinity proteomics to identify extracellular HBPs in pancreas and plasma of normal mice and in a caerulein mouse model of AP. Many new extracellular HBPs (360) were discovered in the pancreas, taking the total number of HBPs known to 786. Extracellular pancreas HBPs form highly interconnected protein-protein interaction networks in both normal pancreas (NP) and AP. Thus, HBPs represent an important set of extracellular proteins with significant regulatory potential in the pancreas. HBPs in NP are associated with biological functions such as molecular transport and cellular movement that underlie pancreatic homeostasis. However, in AP HBPs are associated with additional inflammatory processes such as acute phase response signalling, complement activation and mitochondrial dysfunction, which has a central role in the development of AP. Plasma HBPs in AP included known AP biomarkers such as serum amyloid A, as well as emerging targets such as histone H2A. Other HBPs such as alpha 2-HS glycoprotein (AHSG) and histidine-rich glycoprotein (HRG) need further investigation for potential applications in the management of AP. Pancreas HBPs are extracellular and so easily accessible and are potential drug targets in AP, whereas plasma HBPs represent potential biomarkers for AP. Thus, their identification paves the way to determine which HBPs may have potential applications in the management of AP.

## Introduction

The pancreas develops from endodermal cells in the foregut and has important exocrine and endocrine functions (1). Acute pancreatitis (AP) is acute inflammation of the pancreas and is mainly caused by gallstones and alcohol (2). It is a leading gastrointestinal cause of hospitalization and has significant quality of life implications for the patient and cost implications for health systems (3). Although most episodes of AP are mild and self-limiting, the severe form of the disease, accompanied by a systemic inflammatory response syndrome and multi-organ failure, is associated with a high mortality. At a cellular level disruption of calcium signaling and mitochondrial dysfunction have been implicated in the pathogenesis of AP (4, 5).

The clinical characteristics of AP suggest that an important molecular component is change in cell communication within the pancreas and, in severe AP, systemically between the pancreas and other organs. Cell communication occurs between cells through the medium of the extracellular matrix, with cell receptors responsible for generating the cellular response. The importance of extracellular proteins in mediating communication between cells in multicellular organisms is underscored by the demonstration that the increase in multicellular organism complexity is accompanied by an expansion and increase in the complexity of extracellular proteins (6). A key non-protein component of the extracellular space is the glycosaminoglycan heparan sulfate (HS), because it binds and regulates the activity of a large number of extracellular proteins involved in cell communication ((7); reviewed in (8); (9)).

HS is formed of linear repeats of a disaccharide of 1,4 linked uronic acid (α-L-iduronate, IdoA, or β-D-glucuronate, GlcA) and α-D-N-acetylglucosamine (GlcNAc), with variable O-sulfation of C2 on the uronic acid, of C3 and C6 on the glucosamine; the glucosamine being either N-acetylated or N-sulfated (8, 9). These modifications are hierarchical (9, 10), resulting in chains with a distinct domain structure (11); heparin-binding proteins (HBPs) bind to the sulfated domains and their flanking transition domains. HS chains are attached to proteins to form proteoglycans (HSPGs), with the core proteins serving in part to direct chains to particular extracellular locations (9). Heparin is often used as a proxy for the sulfated domains of HS, though it is more homogenous and sulfated than these (8).

Many individual studies have demonstrated a variety of mechanisms whereby binding to HS regulates the function of extracellular proteins, but only recently has this been analysed on a global scale (7). The latter work demonstrated that of the extracellular proteins, the HBPs form a highly interconnected functional network through protein-protein interactions that are linked to physiological and pathological processes in more complex multicellular organisms. Along with the numerous studies on individual proteins, the HBPs have been demonstrated to be important signalling molecules in the cellular microenvironment that influence fundamental biological processes in development, homeostasis and disease (7, 9, 12).

We previously used mRNA expression data as a proxy for protein to test the hypothesis that HBPs, which are demonstrably functionally important at a genome-wide level (7), are equally important in the context of a single organ, the pancreas, and its associated digestive diseases (13). This work showed that HBPs are indeed a key subclass of extracellular proteins, whose pattern of expression differs in the healthy and diseased pancreas. However, this work has compromises, since mRNA expression does not always correlate with protein expression (14–16). Moreover, the use of mRNA assumes that the existing set of 435 HBPs (7) is reasonably representative of those expressed in pancreas and that this organ does not express novel HBPs. From the perspective of pancreatic diseases, HBPs, because they bind to heparin and are extracellular, represent easily accessible new targets for the development of drugs. However, from a biomarker perspective, sampling pancreas HBPs would be invasive.

We have, therefore, undertaken a proteomic analysis of HBPs in the pancreas and plasma of normal mouse and in the caerulein-induced mouse model of AP. We identified 360 extracellular HBPs in pancreas that were not previously known to bind the polysaccharide. Analysis of the protein-protein interactions of the extracellular pancreas and plasma HBPs demonstrates that they are highly interconnected in NP and AP. Label-free quantification by mass spectrometry identified a number of HBPs, including some known biomarkers of AP, such as carboxypeptidases (CPB1 and CPB2) and pancreatic amylase (AMY2A) (17), which were overexpressed in AP as compared to NP. Isolation of plasma HBPs by heparin affinity chromatography resulted in a reduction of the high abundance protein albumin, so that it would not interfere with mass spectrometry. Of the 151 plasma HBPs isolated, some were known biomarkers of AP, such as serum amyloid A (SAA) while others were emerging novel markers and drug targets in AP such as histone H2A. These may provide a route to applications in the management of AP.

## Methods

### HBPs from pancreas

#### Pancreatic murine models

CD1 mice were used in all experiments, as they are genetically heterogeneous and more likely to represent the human population (18). Pancreases were obtained from 6-8 week old male adult CD1 mice (weight range, 24-30 g) with normal pancreas (NP) or experimental acute pancreatitis (AP). In order to induce experimental acute pancreatitis, CD1 mice were fasted for 12 hours before each experiment, following which they were administered seven hourly intraperitoneal injections of caerulein (50 μg/kg; Sigma-Aldrich, Gillingham UK) dissolved in 0.9% (w/v) saline (Braun Medical, Aylesbury, England). Pancreatitis was confirmed in the experimental mice 24 hours after the first intraperitoneal injection. All mice (NP and AP) were euthanized by cervical dislocation. Studies were conducted in compliance with UK Home Office regulations (PL4003320), and were approved by the Institutional ethical review Committee of the University of Liverpool. A blood sample was collected for serum amylase determination. Serum amylase activity was tested in the Clinical Biochemistry Department in Royal Liverpool University Hospital. A sliver of pancreatic tissue from each of the pancreases was fixed in formalin for H&E staining and histological examination. The pancreases were removed, weighed, pooled together and placed in buffer H (10 mM HEPES pH 7.5, 5 mM MgCl_2_, 25 mM KCl, 0.25 M sucrose supplemented with Complete protease inhibitors cocktail, Roche Products Ltd, Welwyn Garden City, UK) at 4 °C. Complete protease inhibitor was used in all experiments as it has been shown to be particularly effective in experiments involving pancreatic tissue (19). Sixteen CD1 mice were used for each HBP isolation experiment. Each experiment was performed three times for NP and for AP.

#### Isolation of a plasma membrane enriched fraction

The isolation of the plasma membrane from murine pancreases was performed as described (7), with minor modifications. All the steps were performed on ice or at 4 °C. Briefly, mouse pancreases were minced and homogenised with buffer H using a 30 mL Potter-Elvehjem homogeniser (30-40 strokes). Subcellular fractionation was performed using sequential steps of centrifugation. The homogenate was centrifuged for 20 min at 1,000 g in a Sorvall centrifuge (SS-34 rotor, DuPont UK, Stevenage, UK). The pellet was resuspended with buffer H, homogenised and centrifuged again. The two supernatants were then combined (S1) and centrifuged for 20 min at 25,000 g. This supernatant was transferred to a fresh tube and the centrifugation repeated. The final supernatant (S2) was centrifuged for 45 min at 135,000 g in a Sorvall Ultra Pro 80 ultracentrifuge (T.865.1 rotor) to produce a microsomal pellet, which was washed with 8 mL of buffer H (W) by resuspension and centrifuged as above.

The final microsomal pellet was resuspended by homogenisation in a Dounce homogeniser (5 mL) with 4 mL of 1.55 M sucrose in buffer H and placed on a 2 mL 2 M sucrose cushion in a swing-bucket ultracentrifuge tube. It was then overlaid with 2.5 mL 1.33 M, 2 mL 1.2 M, 2 mL 1.1 M, 1 mL 0.77 M and 1 mL 0.25 M sucrose, all in buffer H. The sucrose gradient was centrifuged for 16 h at 116,000 g in a Sorvall Ultra Pro 80 ultracentrifuge (AH-629 rotor). Purdenz™ (GENTAUR Ltd, London, UK) was pumped into the bottom of the tube using a 20 mL syringe to collect 1 mL fractions (F1-F12) from the top of the sucrose gradient. The first eleven fractions (F1-F11) were diluted eight times with buffer H and centrifuged for 45 min at 135,000 g. The last fraction was discarded. The pellets thus obtained (P2-P11) were resuspended with 400 μL 2 % (v/v) Triton X-100 (Sigma Aldrich) in phosphate-buffered saline (PBS; 10 mM Na_2_HPO_4_, 1.8 mM KH_2_PO_4_, 137 mM NaCl and 2.7 mM KCl, pH 7.4) using a Dounce homogeniser (1 mL). Protein concentration was measured by the bicinchoninic acid (BCA) protein assay (Pierce, Thermo Fisher Scientific, Northumberland, UK) in 1:20 dilutions of the resuspended pellets (final Triton X-100 concentration: 0.1 % (v/v)). Equal amounts of the resuspended pellets P1-P11 were analysed by SDS-PAGE and by western blot, using a polyclonal antibody to caveolin1 (sc-890, Santa Cruz Biotechnology Inc., Insight Biotechnology, Wembley, UK). After gel electrophoresis, samples were transferred to Hybond™ nitrocellulose membrane (GE Healthcare Life Sciences, Buckinghamshire, UK) using a wet system (Mini Trans-Blot Cell, Bio-Rad Laboratories Ltd Hemel Hempstead, Hertfordshire, UK) for 1 h at 100 V. Membranes were blocked with Tris-buffered saline (TBS) (20 mM Tris-Cl, 150 mM NaCl, pH 7.5) supplemented with 10 % (w/v) skimmed milk powder for 1 h at room temperature. The membranes were then incubated in TBST-1% (w/v) milk (TBS supplemented with 0.1 % (v/v) Tween 20 (TBST) and 1 % (w/v) skimmed milk powder) with anti-caveolin1 (1:200) overnight at 4 °C. After three 5 min washes with TBST, the membranes were incubated in TBST-5 % w/v milk (TBST supplemented with 5 % (w/v) skimmed milk powder) with anti-rabbit-HRP secondary antibody (A0545, Sigma Aldrich) (1:5,000) for 1 h at room temperature. After at least three 5 min washes with TBST, the membranes were developed using the SuperSignal West Pico Chemiluminescent substrate (Pierce, Thermo Fisher Scientific, Northumberland, UK).

#### Heparin affinity chromatography

Membrane pellets P2-P4 were selected according to their sedimentation profile and caveolin-1 immunoreactivity. The resuspended pellets were pooled, adjusted to 0.15 M NaCl and 1 % (v/v) Triton X-100 and were applied to a 1 mL Hi-Trap heparin column (GE Healthcare Life Sciences) equilibrated with a modified phosphate-buffered saline (PBS), buffer WL (0.15 M NaCl, 13.7 mM Na_2_HPO_4_, 6.3 mM Na_2_PO_4_, 0.1 % (v/v) Triton X-100, pH 7.2). After loading, the column was extensively washed with buffer WL until the absorbance at 280 nm reached the baseline. Bound proteins were then eluted with a single column volume of buffer E (2 M NaCl, 13.7 mM Na_2_HPO_4_, 6.3 mM Na_2_HPO_4_, 0. 1 % (v/v) Triton X-100, pH 7.2). Protein concentration was measured by the BCA protein assay.

#### Sample preparation for mass spectrometry

Trichloroacetic acid (TCA, 70% w/v) was added to an equal volume of the heparin-bound fraction and kept at −20°C for 1 hour, followed by centrifugation at 14,000 rpm for 10 minutes. After removing the supernatant fraction carefully with a glass pipette, the pellet was washed 5 times with 5% (w/v) TCA. The pellet was freeze-dried overnight, washed subsequently with 0.5 mL diethyl ether to remove the excess TCA and centrifuged at 5,000 rpm. The supernatant was removed and the diethyl ether wash was repeated two more times. The final pellet was left to dry in the fume hood. The ether-washed, TCA-precipitated pellets were resolublised in either 200 μL (normal pancreas (NP)) or 600 μL (acute pancreas (AP)) 50 mM ammonium bicarbonate, 0.05 % (w/v) Rapigest (Waters, Manchester) and shaken at 550 rpm for 10 min at 80 °C. The sample was then reduced (addition of 10 μL (NP) or 30 μL (AP) of 60 mM DTT and incubation at 60 °C for 10 min) and alkylated (addition of 10 μL (NP) or 30 μL (AP) of 180 mM iodoacetamide and incubation at room temperature for 30 minutes in the dark). Trypsin (Sigma, Poole, UK, proteomics grade) was reconstituted in 50 mM acetic acid to a concentration of 0.2 μg/μL and 10 μL (NP) or 30 μL (AP) added to the sample followed by overnight incubation at 37 °C. The digestion was terminated and RapiGest™ was removed by acidification with the addition of 3 μL (NP) or 9 μL (AP) of 100% v/v trifluoroacetic acid (TFA) and incubation at 37 °C for 45 min) and centrifugation (15,000 x g for 15 min). To check for completeness of digestion each sample was analysed pre- and post-acidification by SDS-PAGE.

#### Mass spectrometry data acquisition and analysis

For LC-MS/MS analysis each digest was diluted to 250 ng/μL with 97/3/0.1 % (v/v) water/acetonitrile/formic acid and mixed 2:1 with a protein digest standard (50 fmol/μL yeast alcohol dehydrogenase, Mass PREP™ Digestion Standard, Waters). A 3 μL injection of this mixture, corresponding to 500 ng of sample and 50 fmol of standard was analysed using an Ultimate 3000 RSLC™ nano system (Thermo Scientific, Hemel Hempstead, U.K.) coupled to a QExactive™ mass spectrometer (Thermo Scientific). The sample was loaded onto the trapping column (Thermo Scientific, PepMap100, C18, 300 μm X 5 mm), using partial loop injection, for 7 min at a flow rate of 4 μL/min with 0.1 % (v/v) TFA. The sample was resolved on the analytical column (Easy-Spray C18 75 μm x 500 mm 2 μm column) using a gradient of 97 % A (0.1 % formic acid) 3 % B (99.9 % acetonitrile, 0.1 % formic acid) to 60 % A 40 % B (all v/v) over 90 minutes at a flow rate of 300 nL/min. The data-dependent program used for data acquisition consisted of a 70,000 resolution full-scan MS scan (automatic gain control (AGC) set to 1e6 ions with a maximum fill time of 250 ms) and the 10 most abundant peaks were selected for MS/MS using a 17,000 resolution scan (AGC set to 5e4 ions with a maximum fill time of 250 ms) with an ion selection window of 3 *m/z* and a normalised collision energy of 30. To avoid repeated selection of peptides for MS/MS, the program used a 30 second dynamic exclusion window. The data were processed with Progenesis QI (version 2 Nonlinear Dynamics, Newcastle upon Tyne, UK). Samples were aligned according to retention time using a combination of manual and automatic alignment. Default peak picking parameters were applied and features with charges from 1^+^ to 4^+^ and with three or more isotope peaks were retained. Database searching was performed using Mascot (Matrix Science, London, UK). A Mascot Generic File, created by Progenesis QI, was searched against the reviewed entries of the reference proteome set of *M. musculus* from Uniprot (19/02/2014, 43238 sequences) with the sequence of yeast alcohol dehydrogenase (UniProt: P00330) added. A fixed carbamidomethyl modification for cysteine and variable oxidation modification for methionine were specified. A precursor mass tolerance of 10 ppm and a fragment ion mass tolerance of 0.01 Da were applied. The results were then filtered to obtain a peptide false discovery rate (FDR) of 1 % and further refinement of two peptides per protein was applied. Label-free quantification was performed following the “Top3” methodology (20) by the spiking of the sample prior to analysis with the internal standard of 50 fmol yeast alcohol dehydrogenase digest (Uniprot P00330, Waters). Proteins were annotated as differentially expressed if they achieved a FDR corrected q value of 1 %. Additionally, an adjusted threshold p value of less than 0.001 following the Bonferroni correction was used to identify the HBPs as potential biomarkers (21).

#### Identification of extracellular HBPs in NP and AP

Extracellular proteins were identified using a combination of bioinformatics tools. SignalP 4.1, which predicts the presence of a secretory signal peptide, was used to identify extracellular proteins (22) with Phobius, which is a combined transmembrane topology and signal peptide prediction tool, to obtain a wider coverage for extracellular protein identification (23). A third tool, SecretomeP 2.0, was used to produce *ab initio* predictions of protein secretion not based on a secretory signal peptide (24). A fourth tool QIAGEN’s Ingenuity Pathway Analysis (IPA^®^, QIAGEN Redwood City, www.qiagen.com/ingenuity), was used to identify extracellular and plasma membrane proteins. The HBPs that were not identified using SignalP, Phobius, or SecretomeP, but which were identified using IPA were further investigated using a manual approach. In addition, candidate HBPs identified by SignalP, Phobius or SecretomeP that were classified as not being extracellular using IPA were further examined using a manual approach. Each candidate HBP was examined using UniProtKB (25) for the presence of an extracellular signature. The outputs of the various approaches were merged to obtain the complete list of HBPs in NP and AP.

#### Interactions and construction of networks of HBPs

Interactions between HBPs were obtained from the online database resource ‘Search Tool for the Retrieval of Interacting Genes’ (STRING). STRING 10.5 is a database of known and predicted functional interactions and served as a ‘one-stop’ comprehensive resource for network creation and analysis (26). The interactions in STRING are provided with a probabilistic confidence score that is an estimate of how likely an interaction describes a functional linkage between two proteins. A higher score indicating a higher confidence is attributed when more than one type of information supports a given association. Only interactions with a high confidence score (0.70 and above) were used to build HBP networks in NP and AP. The resulting networks were termed ‘protein interactomes’. A network is deemed to be well connected if its average clustering coefficient is significantly higher than that of its corresponding random networks (27).

#### Functional analyses of extracellular pancreas HBPs in NP and AP

*Canonical pathways and bio-functions analyses* - Canonical pathways are well-characterized signalling pathways that have been curated from original data. Bio-functions are molecular and cellular functions that play important roles in homeostasis in health and disease. The significance of the association between the datasets and the canonical pathway/bio-function was measured by calculating the p-value using Fisher’s exact test to determine the probability of the association between the HBPs in the dataset and the canonical pathway/bio-function. Canonical pathways and bio-function analyses are useful tools for the identification of potential biomarkers and drug targets. These were undertaken using IPA. They also help identify biological pathways that play important roles in homeostasis and complex diseases.

### Plasma HBPs

This method is deposited at protocols.io (***dx.doi.org/10.17504/protocols.io.wdtfa6n***).

#### Sample collection

Blood obtained from 6-8 week old male adult CD1 mice (weight range, 24-30 g) with normal pancreas (NP) or experimental acute pancreatitis (AP), induced as described above, was collected in 0.38 % (w/v) sodium citrate (Sigma-Aldrich, Gillingham, UK) and centrifuged at 5,000 g at 4 °C for 15 min (28). Supernatant plasma was extracted and frozen at −20 °C.

#### Heparin affinity chromatography

The frozen murine plasma was defrosted on ice and centrifuged at 16,100 g for 10 min, at 4°C. The resulting supernatant was diluted (1:8) in 75 mM NaCl, 6.85 mM Na_2_HPO_4_, 3.15 mM NaH_2_PO_4_, pH 7.2. This method was effective in reducing the high abundance protein, albumin, down to levels that would not affect the MS analysis of plasma HBPs. The diluted supernatant was centrifuged at 5,000 g for 5 min and the resulting supernatant was then applied on a 1 mL Hi-Trap heparin column (GE Healthecare Life Sciences), equilibrated in modified PBS. The heparin column was subsequently washed extensively with 50 mL of modified PBS (150 mM NaCl, 13.7 mM Na_2_HPO_4_, 6.3 mM NaH_2_PO_4_, pH 7.2). The heparin-bound fraction was then eluted with 2 M NaCl in modified PBS.

#### Protein adsorption using StrataClean for mass spectrometry

In order to eliminate the high concentration of electrolytes (2.15 M NaCl) in the eluate, so as not to influence MS analysis, StrataClean™ beads (Agilent Technologies, UK) were used to adsorb proteins (29). Briefly, 100 μg of the heparin-bound fraction was mixed with 30 μL StrataClean slurry, vortexed for 2 min. and centrifuged at 2,000 g for 2 min at 4 °C. The supernatant was removed carefully and discarded. The pellet was first washed with PBS and then further washed twice with H_2_O.

#### Sample preparation for mass spectrometry

In the case of plasma samples, the StrataClean™ beads were resuspended in 80 μL of 25 mM ammonium bicarbonate and 5 μL of 1 % (w/v) Rapigest (Waters, Manchester, UK) added and the samples shaken at 450 rpm for 10 min at 80 °C. Samples were reduced by the addition of 5 μL of 60 mM Dithiothreitol (DTT) and incubated at 60 °C for 10 min and alkylated (addition of 5 μL of 180 mM iodoacetamide and incubation at room temperature for 30 min in the dark). Trypsin (Sigma, Poole, UK, proteomics grade) was reconstituted in 50 mM acetic acid to a concentration of 0.2 μg/μL and 5 μL (1 μg) added to the sample followed by overnight incubation at 37 °C. Samples were mixed on a rotating mixer in overnight at 37 °C. The following day the digestion was terminated and Rapigest removed by acidification with TFA (1 μL) and incubation at 37 °C for 45 min. Samples were centrifuged at 17,200 g for 30 min and the clarified supernatants transferred to tubes.

#### Mass spectrometry data acquisition and analysis

Data acquisition and analysis were undertaken similarly with some modifications. Briefly, for LC-MS/MS analysis each digest was mixed with an equal volume of a protein digest standard (50 fmol/μL yeast enolase, Mass PREP™ Digestion Standard, Waters). A 2 μL injection of this mixture, corresponding to 50 fmol of standard was analysed using an Ultimate 3,000 RSLC™ nano system (Thermo Scientific, Hemel Hempstead, U.K.) coupled to a QExactive™ mass spectrometer (Thermo Scientific). The sample was loaded onto the trapping column (Thermo Scientific, PepMap100, C18, 300 μm x 5 mm), using partial loop injection, for 7 min at a flow rate of 9 μL/min with 0.1 % (v/v) TFA/2 % (v/v) acetonitrile. The sample was resolved on the analytical column (Easy-Spray C18 75 μm x 500 mm 2 μm column) using a gradient of 97 % A (0.1 % formic acid) 3 % B (99.9 % acetonitrile, 0.1 % formic acid) to 60 % A 40 % B (all v/v) over 90 min at a flow rate of 300nL/min. The data-dependent program used for data acquisition consisted of a 70,000 resolution full-scan MS scan (automatic gain control (AGC) set to 1e6 ions with a maximum fill time of 250 ms) and the 10 most abundant peaks were selected for MS/MS using a 35,000 resolution scan (AGC set to 1e5 ions with a maximum fill time of 100 ms) with an ion selection window of 2 m/z and a normalised collision energy of 29. To avoid repeated selection of peptides for MS/MS, the program used a 20 second dynamic exclusion window. The data were processed with Progenesis QI (version 3 Waters, Newcastle upon Tyne, UK). Samples were automatically aligned according to retention time. Default peak picking parameters were applied and features with charges from 2+ to 7+ and were retained. Database searching was performed using Mascot (Matrix Science, London, UK). A Mascot Generic File, created by Progenesis QI, was searched against the reviewed entries of the reference proteome set of M. musculus from Uniprot (23/06/2015, 16,711 sequences) with the sequence of yeast enolase (UniProt: P00924) added. A fixed carbamidomethyl modification for cysteine and variable oxidation modification for methionine were specified. A precursor mass tolerance of 10 ppm and a fragment ion mass tolerance of 0.01 Da were applied. The results were then filtered to obtain a peptide false discovery rate of 1 %. Label-free quantification was performed using the “Top3” methodology (20) by the spiking of the sample prior to analysis with the internal standard of 50 fmol yeast enolase (UniProt: P00924, Waters).

## Results

### Isolation of pancreas HBPs from plasma membrane enriched fractions from normal pancreas and a murine model of acute pancreatitis

Each extracellular pancreas HBP isolation experiment was performed three times using sixteen pancreases from control (NP) and AP mice. Acute pancreatitis was induced in animals by injection of caerulein. Histological analysis of samples of these pancreases demonstrated the classic features of normal pancreas with preserved acinar pattern (Fig. 1A) and those associated with AP, namely marked oedema, vacuolisation, neutrophil infiltration in the ductal margins and parenchyma of the pancreas, with focal acinar cell necrosis (Fig. 1B). Serum amylase increased ~8- to 10-fold in AP compared to NP (Figs. 1C, D). These data are consistent with successful induction of AP (30).

**Figure 1:**
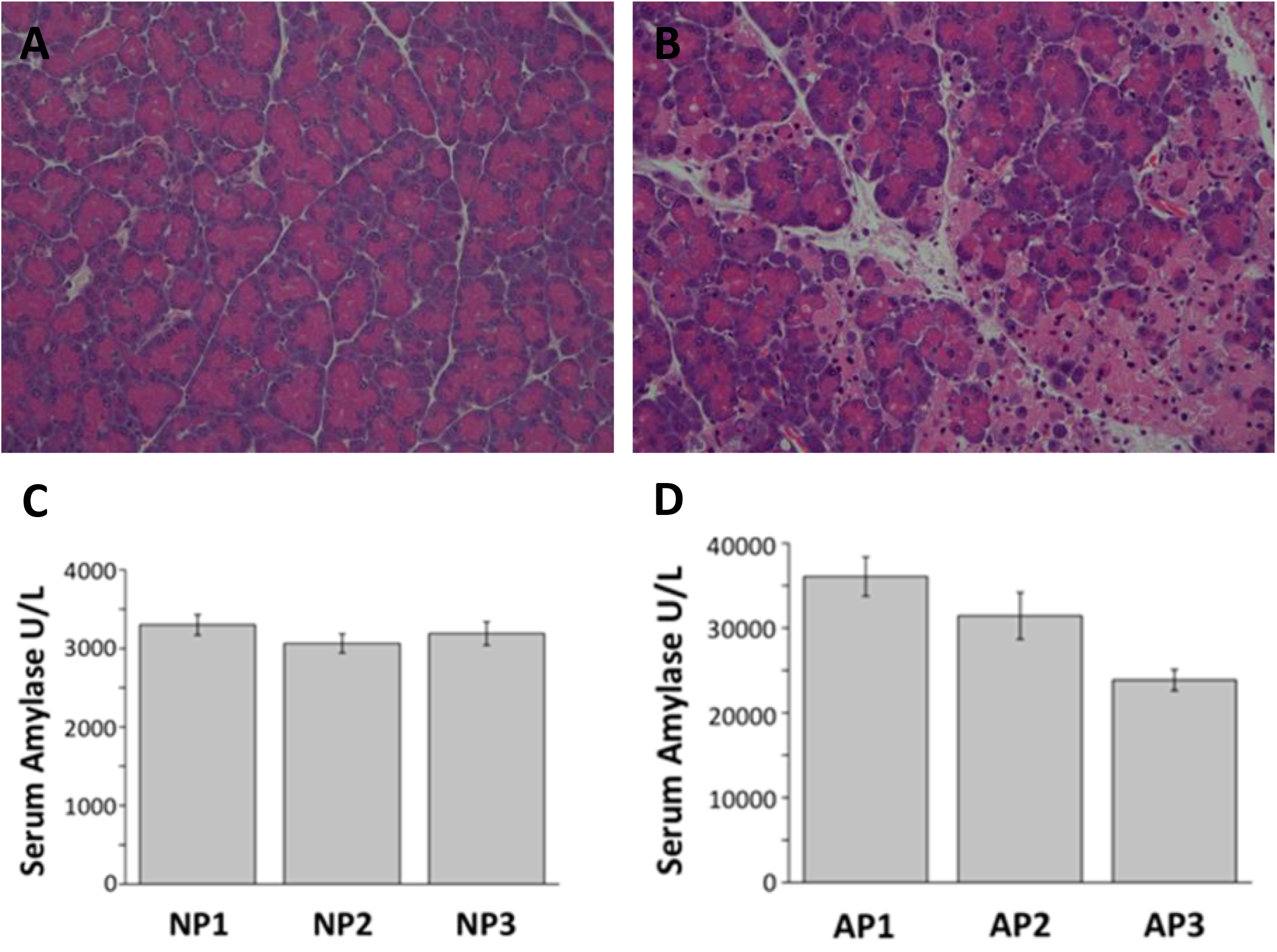
Normal pancreas (NP) and caerulein-induced acute pancreatitis (AP). Representative images of H&E stained histology slides of A) NP with intact pancreas architecture and B) AP showing marked oedema, inflammatory cell infiltration and acinar cell necrosis. Mean serum amylase levels in (C) NP and (D) AP in each experiment consisting of 16 individuals.

Differential centrifugation (31, 32) of pancreas homogenates produced a series of cellular sub-fractions, including the final microsomal pellet (lane Mc underlined red, Figs. 2A, 2B), which was floated on a sucrose gradient (0.77-1.55 M) to separate its individual constituents (Figs. 2C, 2D). The gradient was harvested into 12 fractions, with the final fraction being discarded. Eleven fractions were assessed by Western blot for the plasma membrane marker caveolin-1 (Figs. 2C, 2D). The caveolin-1 content was inversely correlated with the equilibrium density of the sucrose fractions, consistent with their identification as plasma membranes, which possess lower density than other microsomal membranes. Fractions 2 to 4 (Figs. 2C, 2D underlined red), which had the strongest caveolin-1 immunoreactivity, were pooled and selected as the plasma membrane enriched fraction (PM). In addition to membrane-resident proteins, which are often underrepresented in proteomic analyses, this fraction would also contain intra- and extracellular membrane-associated proteins, as well as proteins associated with the pericellular matrix.

**Figure 2:**
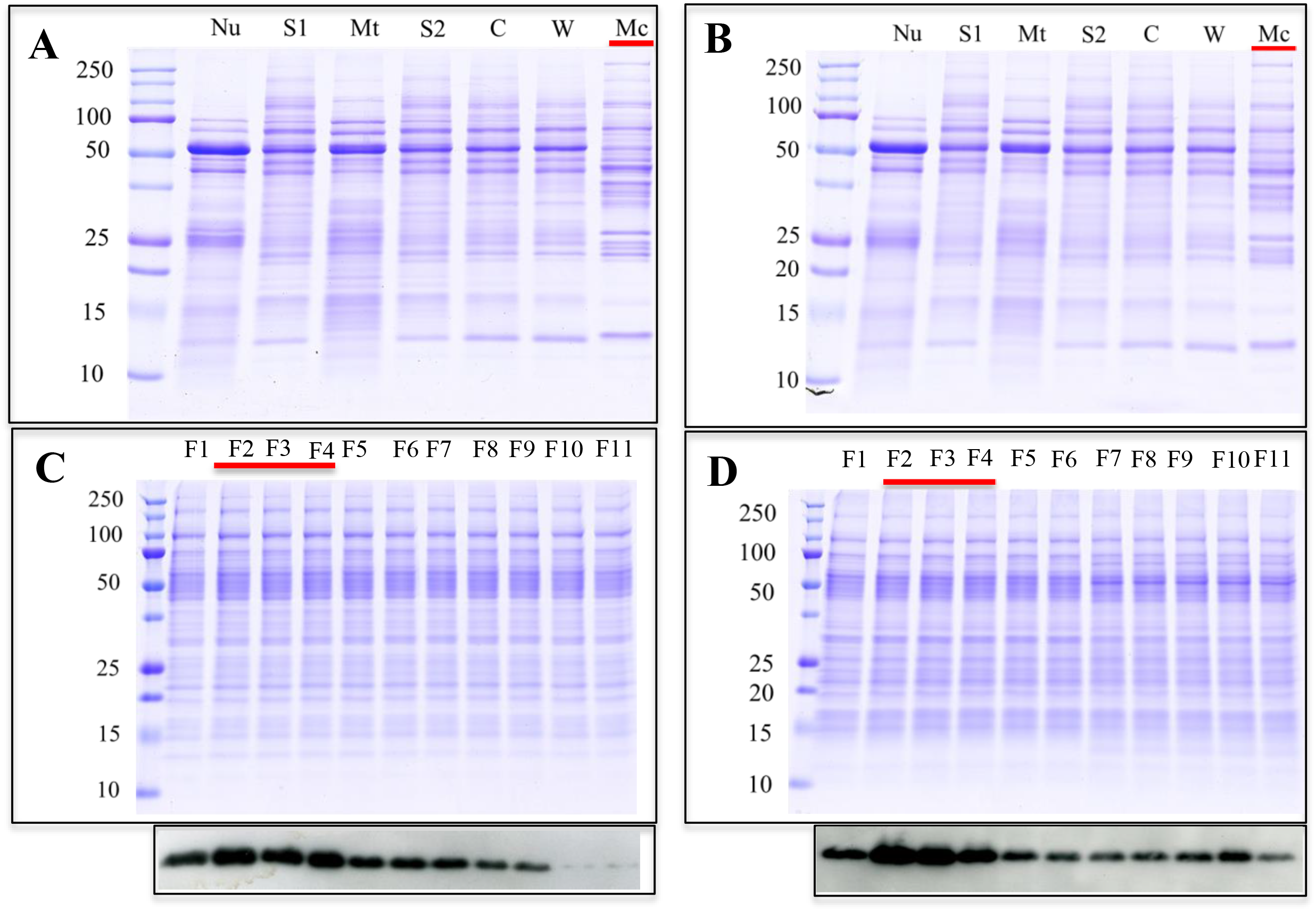
Preparation of a plasma membrane enriched fraction. Coomassie-stained SDS-PAGE gel of (A) NP and (B) AP samples obtained during homogenisation and fractionation by sequential steps of centrifugation. Nu = nuclear pellet; S1 = post-nuclear supernatant; Mt = mitochondrial pellet; S2 = post-mitochondrial supernatant; C = cytosol (post-microsomal supernatant); W = wash of the microsomal pellet; Mc = microsomal pellet. Coomassie-stained SDS-PAGE and western blot analysis of 10 fractions (F1-F11) from the microsomal pellet after flotation on a sucrose gradient (0.25 - 2 M) in (C) NP and (D) AP. Fractions are ordered depending on their equilibrium density from light (left) to heavy (right). The enrichment of plasma membrane was assessed by western blot using an antibody against caveolin-1, which is a specific plasma membrane marker. Full Western blots in Fig. S1.

Fractions 2 to 4 were solubilised in 1 % (v/v) Triton-X-100 and subjected to heparin affinity chromatography. After extensively washing the heparin column with PBS with 0.1 % (v/v) Triton-X-100, proteins that remained bound were deemed to have a sufficiently strong interaction with the polysaccharide to be considered as HBPs. In contrast to previous work (7), here a single elution at 2 M NaCl with 0.1 % (v/v) Triton X-100 was used to recover heparin-binding proteins. The rationale was that the concentration of NaCl required for elution from heparin does not necessarily correlate with affinity for HS (33, 34), so fractionating the HBPs does not necessarily yield useful information.

#### MS analysis of the pancreas HBPs

Proteins in the 2 M NaCl eluates were precipitated with TCA to remove Triton-X-100, and then digested with trypsin. After ascertaining the optimal loading concentration, the sample order was randomised across biological (three each NP and AP) and technical repeats prior to LC-MS. The LC-MS runs were then individually searched using the MASCOT protein search engine (www.matrixscience.com). The mass spectrometry proteomics data have been uploaded into the PRIDE database (http://www.ebi.ac.uk/pride/archive/) (35), via its partner repository, the ProteomeXchange Consortium (36), with the dataset identifier PXD001950. There was little variation between the technical replicates for the samples, which can be attributed to the high quality of the sample preparation and MS analysis (Fig. 3). Each technical replicate produced between 1500-1900 protein hits at a peptide FDR of 1 % To obtain a fuller coverage, the data were run through Progenesis label-free software. The merged file yielded over 1,900 hits at a peptide FDR of 1 %. Using a two-peptide stringency, these were reduced to 1,602 proteins in NP and 1,866 proteins in AP (Supplementary Tables 1 and 2).

**Figure 3:**
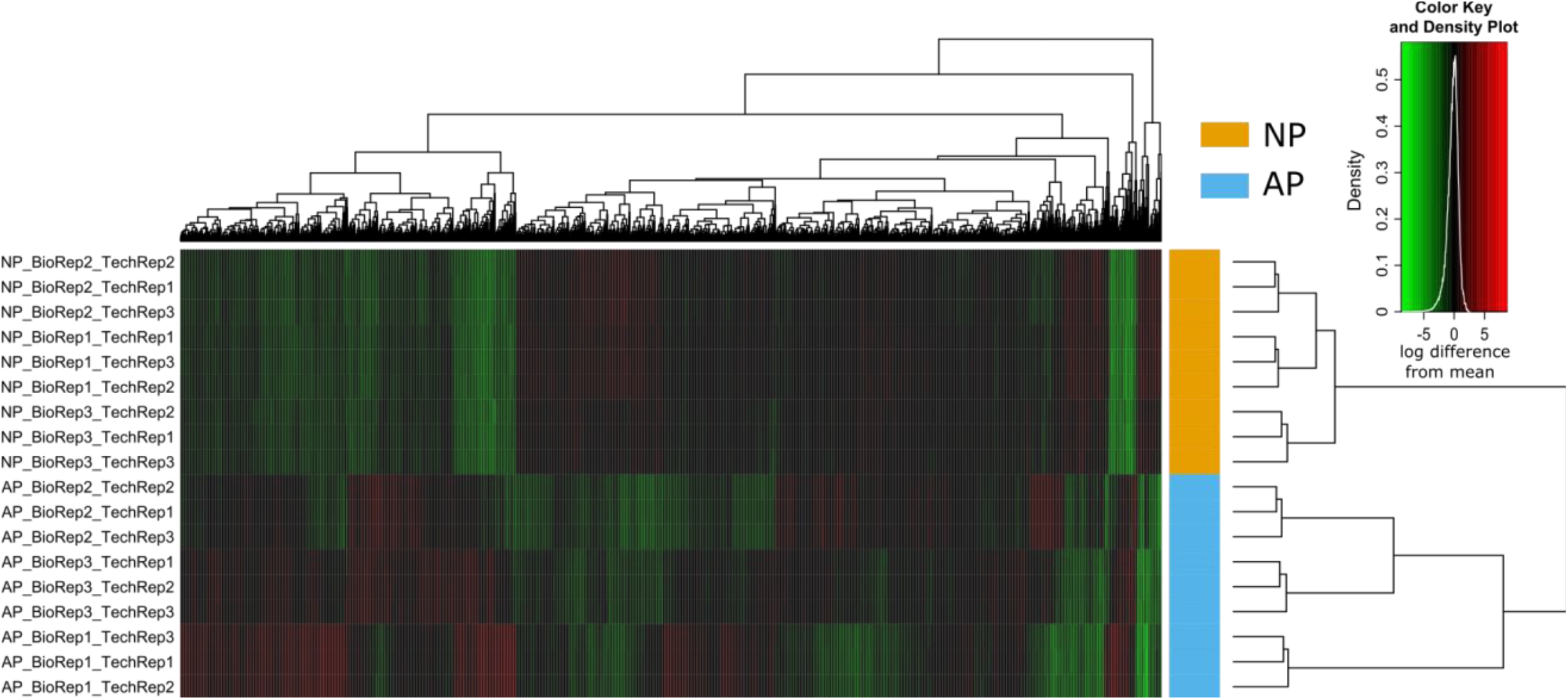
Heat map depicting the variation across the biological and technical replicates in extracellular pancreas HBPs. The rows represent the various biological replicates in normal pancreas (NP) and acute pancreatitis (AP), while the columns represent proteins. Red represents over expression and green represents under expression. Biological replicate number is denoted as “BioRep” and technical replicate number as “TechRep”. Hierarchical clustering was performed on both column data, to cluster the changes in protein expression, and row data, which displays the variation between samples. The stability of the instrument platform is shown in that the lowest branch of the sample variation dendrogram correctly represents the technical replicates of each sample. The next level of the dendrogram correctly separates the biological condition of the sample indicating repeatable protein expression differences between the biological conditions. A higher degree of variability is observed in the AP samples presumably reflecting the systemic effects of AP.

#### Identification of the extracellular HBPs

The 1,602 proteins in NP and 1,866 proteins in AP were then filtered by the bioinformatics pipeline described in “Experimental Procedures” to identify those possessing extracellular (partially or wholly) amino acid sequence. Application of this pipeline required in many instances multiple iterations, because no single approach provided unequivocal evidence for an extracellular location of at least part of the protein sequence. Moreover, a greater confidence was obtained if more than one tool demonstrated extracellular localization. For example, SignalP and Phobius identify proteins with signal peptides, which may be secreted extracellularly, located on the plasma membrane or on an internal cell membrane. Thus secretomeP and IPA provided both a filter for the output of SignalP and Phobius, as well as identifying new candidate extracellular proteins. IPA in particular identified a good many proteins as extracellular that were not identified as such by these other tools. In all cases information on protein localisation and function was examined in UniprotKB. When only one tool, e.g., IPA indicated extracellular localisation, confirmation was sought either through UniprotKB and/or the literature. Filtering the proteins as described above resulted in 320 proteins in NP (Supplementary Table 3) and 345 proteins in AP (Supplementary Table 4) being identified as heparin binding and extracellular. Combining these two sets of proteins yielded 440 HBPs, of which 360 had not been identified previously as heparin binding. With such a substantial number of new HBPs, it was important to analyse their functions and relationships to gain insight into the significance of the pancreas and AP associated HBPs.

#### Label-free quantification

Using the “Top3” methodology, proteins were annotated as differentially expressed between NP and AP if they achieved a FDR corrected q value of 1 % (Supplementary Tables 5 and 6). Introduction of a p value cut off of 0.001, following the Bonferroni correction, resulted in the identification of 79 HBPs that were overexpressed and 48 HBPs that were under expressed significantly in AP, as compared to NP. Known biomarkers of AP, such as carboxypeptidases (CPB1 and CPB2) and pancreatic amylase (AMY2A) (17) were overexpressed in the AP group. The top 20 HBPs with the highest fold change in each group (Tables 1 and 2) may provide further potential biomarkers for AP. Some of the most highly expressed HBPs such as neutrophilic granule protein (NGP) and histidine-rich glycoprotein (HRG) may be representative of the underlying role of innate immunity in early AP (37–39).

**Table 1.**
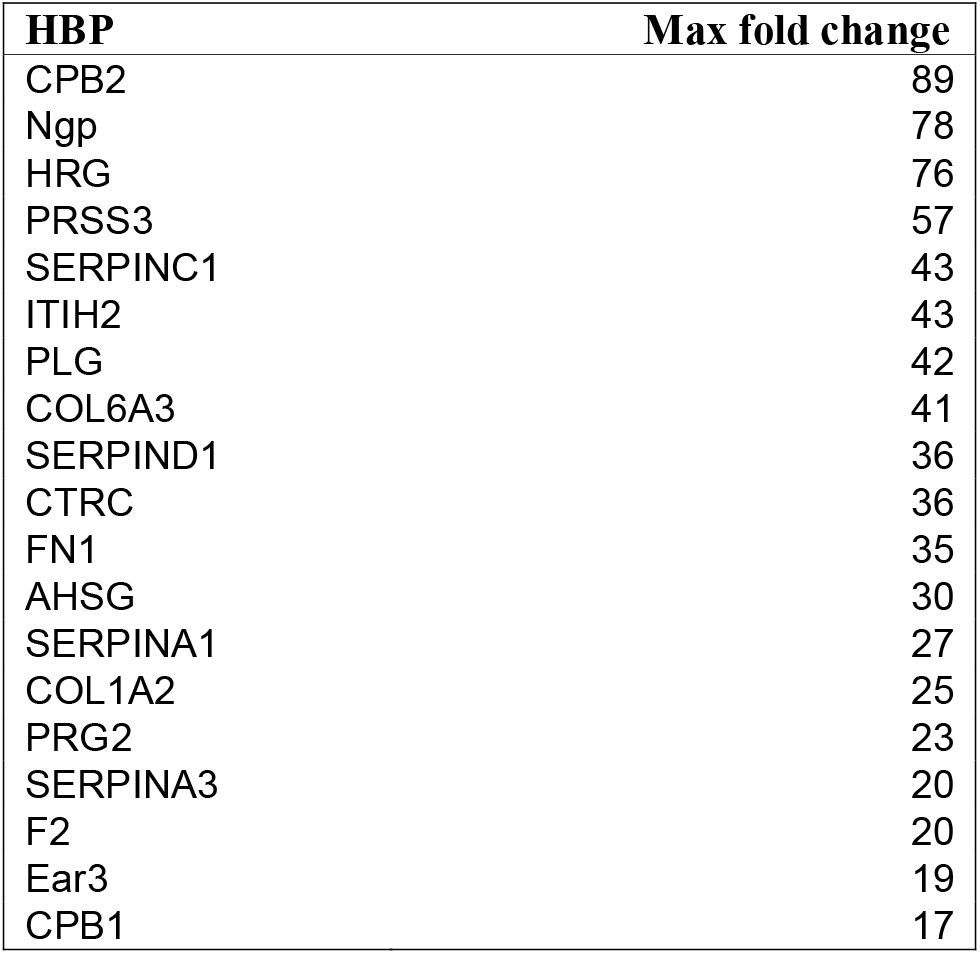
Top 20 extracellular pancreas HBPs overexpressed in AP. The upregulated HBPs were filtered depending on the maximum fold change values. An adjusted threshold p value of less than 0.001 following the Bonferroni correction was used to identify the top HBPs to be validated as potential biomarkers.

| HBP | Max fold change |
| --- | --- |
| CPB2 | 89 |
| Ngp | 78 |
| HRG | 76 |
| PRSS3 | 57 |
| SERPINC1 | 43 |
| ITIH2 | 43 |
| PLG | 42 |
| COL6A3 | 41 |
| SERPIND1 | 36 |
| CTRC | 36 |
| FN1 | 35 |
| AHSG | 30 |
| SERPINA1 | 27 |
| COL1A2 | 25 |
| PRG2 | 23 |
| SERPINA3 | 20 |
| F2 | 20 |
| Ear3 | 19 |
| CPB1 | 17 |

**Table 2.**
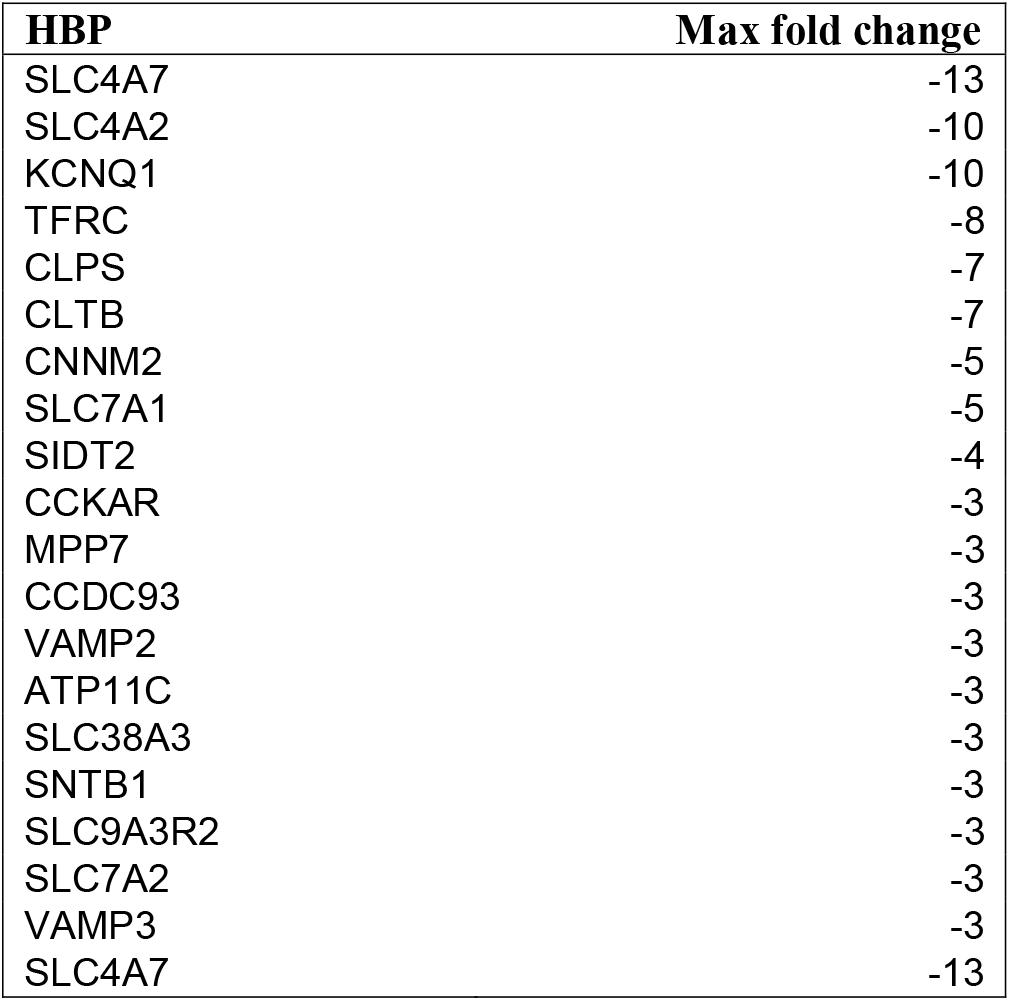
Top 20 extracellular pancreas HBPs underexpressed in AP. The downregulated HBPs were filtered depending on the maximum fold change values. An adjusted threshold p value of less than 0.001 following the Bonferroni correction was used to identify the top HBPs to be validated as potential biomarkers.

| HBP | Max fold change |
| --- | --- |
| SLC4A7 | -13 |
| SLC4A2 | -10 |
| KCNQ1 | -10 |
| TFRC | -8 |
| CLPS | -7 |
| CLTB | -7 |
| CNNM2 | -5 |
| SLC7A1 | -5 |
| SIDT2 | -4 |
| CCKAR | -3 |
| MPP7 | -3 |
| CCDC93 | -3 |
| VAMP2 | -3 |
| ATP11C | -3 |
| SLC38A3 | -3 |
| SNTB1 | -3 |
| SLC9A3R2 | -3 |
| SLC7A2 | -3 |
| VAMP3 | -3 |
| SLC4A7 | -13 |

##### Network construction and analysis of HBPs in the pancreas

The lists of HBPs in NP and AP were used to obtain protein-protein interactions (PPI) from STRING. Only interactions with a high confidence score (0.70 and above) were used to construct extracellular heparin-binding protein interactomes in NP (Fig. 4) and AP (Fig. 5). The topological parameters of the HBP interactomes were obtained from STRING 10.5. The extracellular HBP interactomes of NP and AP have high clustering coefficients (NP = 0.364 and AP = 0.435), which were also significantly higher than those of their corresponding random networks. The PPI enrichment p-values for both extracellular pancreas HBP interactomes were < 1.0e-16. These findings suggest that the HBP interactomes form highly interconnected modules (27, 40) in the extracellular space of the pancreas. Such high connectivity generally reflects a high capacity for regulation. This would mean that the extracellular HBP interactomes are likely to be important in homeostasis of the normal pancreas and the HBPs identified in AP may have key roles in mediating the altered cell communication that is associated with the disease.

**Figure 4:**
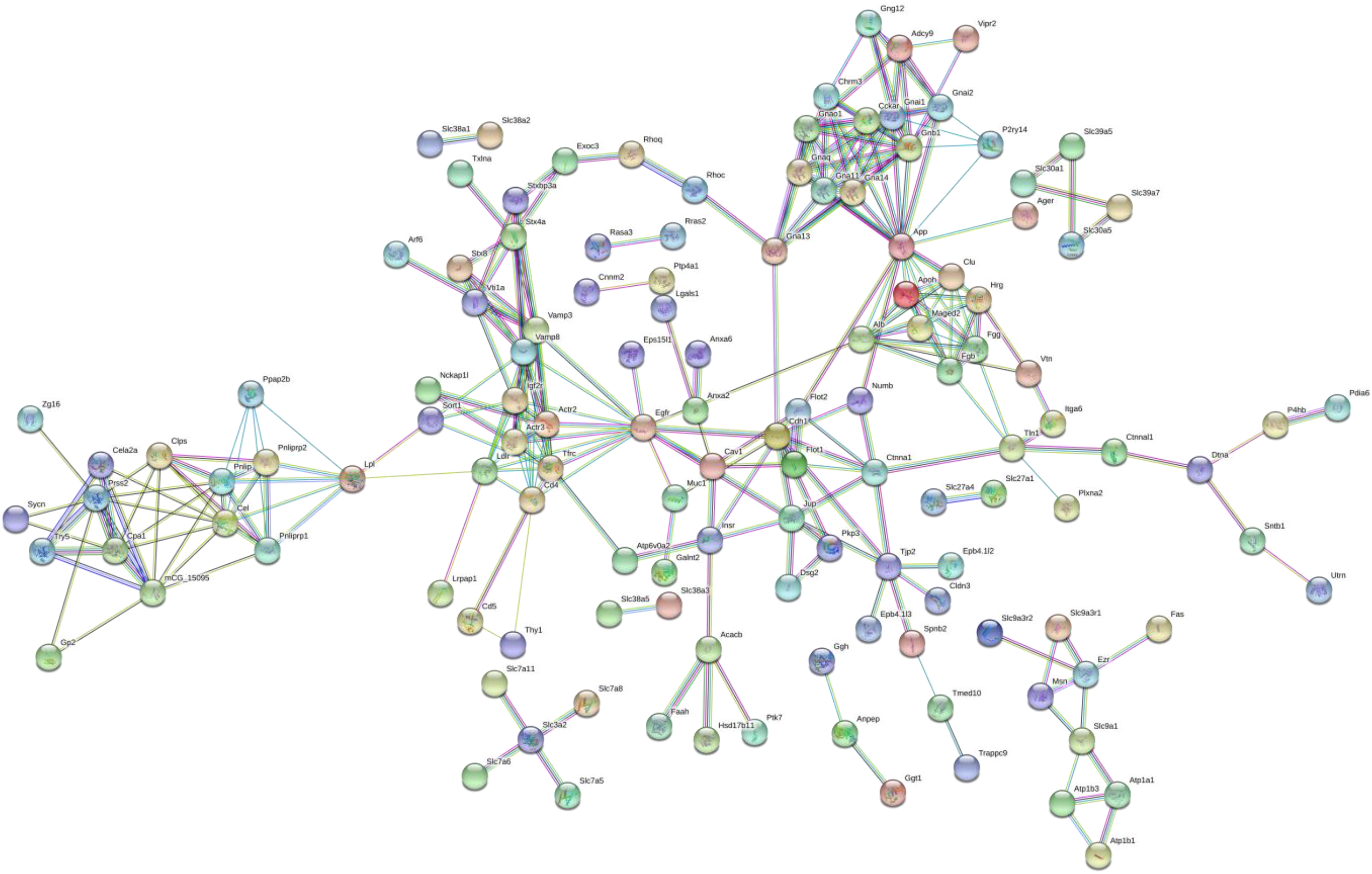
The heparin-binding putative protein interactome in normal pancreas (NP) constructed using STRING 10.5. Nodes or HBPs are connected by protein-protein interactions known as ‘edges’.

**Figure 5:**
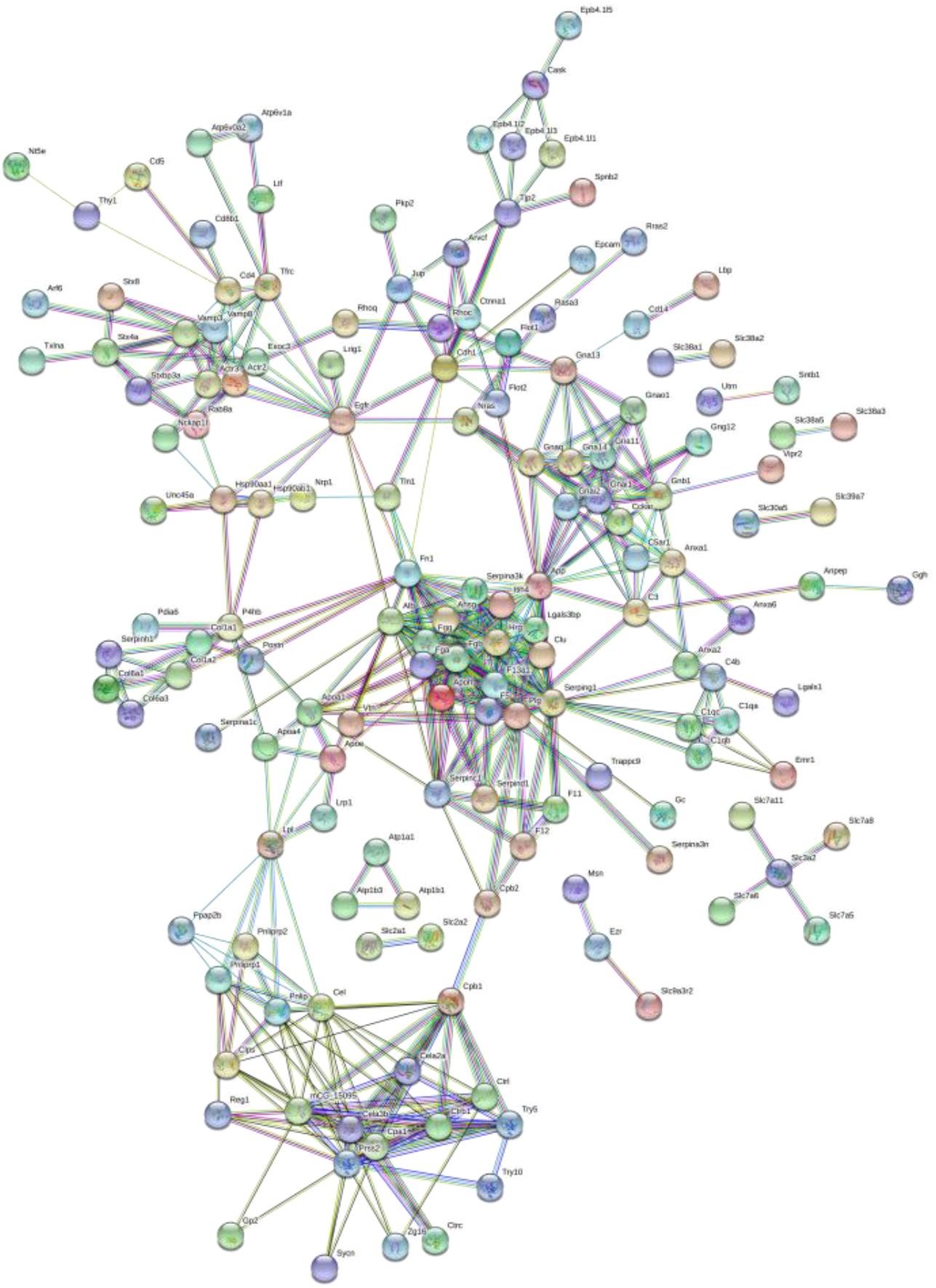
The heparin-binding putative protein interactome in acute pancreatitis (AP) constructed using STRING 10.5. Nodes or HBPs are connected by protein-protein interactions known as ‘edges’.

##### Analysis of Canonical Pathways, Bio-functions of HBPs in NP and AP

The functional context of a particular HBP with respect to the other HBPs will be important, particularly since these proteins are highly interconnected. HBPs in NP and AP clearly have functions important in cell communication and they are enriched in a number of canonical pathways (Tables 3 and 4, Supplementary Tables 7 and 8) underlying homeostasis and complex diseases, both at a systemic and an organ / disease level (NP and AP). For example, the top canonical pathway associated with the NP dataset, ‘Signalling by Rho Family GTPases’ plays an important role in cell adhesion, as do a number of the other highly ranked pathways, ‘RhoGDI Signaling’, ‘Ephrin B Signaling’, ‘Tec Kinase Signaling’, ‘Actin Cytoskeleton Signaling’ and ‘Ephrin Receptor Signaling’. Cell adhesion is essential to the maintenance of tissue architecture and so cell communication appropriate for homeostasis (41). In contrast, the top pathways associated with the HBPs in AP are associated with inflammatory responses. Thus, the top four pathways in AP are ‘Coagulation System’, ‘Acute Phase Response Signalling’, ‘Intrinsic Prothrombin Activation Pathway’ and ‘LXR/RXR Activation’. The presence of pathways linked to cell adhesion further down the list (‘Actin Cytoskeleton Signaling’, ‘RhoGDI Signaling’, ‘Signaling by Rho Family GTPases’, ‘Ephrin Receptor Signaling’, ‘Ephrin B Signaling’ and ‘Integrin Signaling’), would then reflect the predominance of cell motility and a change in tissue architecture, driven by the inflammatory pathways. Analysis of the bio-functions associated with the HBPs in NP (Supplementary Table 9) and AP (Supplementary Table 10) support the conclusions reached from the analysis of canonical pathways.

**Table 3.** Top 20 canonical pathways enriched to extracellular pancreas HBPs in NP using Ingenuity Pathways Analysis. The significance of the association between the datasets and the canonical pathway was measured by calculating the p-value using Fisher’s exact test to determine the probability of the association between the HBPs in the dataset and the canonical pathway.

| Canonical Pathways | -log (p-value) |
| --- | --- |
| Signaling by Rho Family GTPases | 11.1 |
| RhoGDI Signaling | 10.1 |
| CXCR4 Signaling | 9.55 |
| G Beta Gamma Signaling | 8.76 |
| Thrombin Signaling | 8.25 |
| IL-1 Signaling | 8.15 |
| Role of NFAT in Regulation of the Immune Response | 7.98 |
| Tec Kinase Signaling | 7.75 |
| Clathrin-mediated Endocytosis Signaling | 7.67 |
| Caveolar-mediated Endocytosis Signaling | 7.23 |
| Ephrin B Signaling | 7.11 |
| Actin Cytoskeleton Signaling | 6.86 |
| Role of Tissue Factor in Cancer | 6.8 |
| Cardiac Hypertrophy Signaling | 6.51 |
| Integrin Signaling | 6.38 |
| Relaxin Signaling | 6.25 |
| Gαi Signaling | 5.97 |
| Ephrin Receptor Signaling | 5.94 |
| CREB Signaling in Neurons | 5.63 |
| Tight Junction Signaling | 5.45 |

**Table 4.** Top 20 canonical pathways enriched to extracellular pancreas HBPs in experimental AP using Ingenuity Pathways Analysis. The significance of the association between the datasets and the canonical pathway was measured by calculating the p-value using Fisher’s exact test to determine the probability of the association between the HBPs in the dataset and the canonical pathway.

| Canonical Pathways | -log (p-value) |
| --- | --- |
| Coagulation System | 14.9 |
| Acute Phase Response Signaling | 14 |
| Intrinsic Prothrombin Activation Pathway | 13.6 |
| LXR/RXR Activation | 13.4 |
| FXR/RXR Activation | 11.8 |
| Clathrin-mediated Endocytosis Signaling | 11.6 |
| Extrinsic Prothrombin Activation Pathway | 10.1 |
| Complement System | 9.89 |
| Actin Cytoskeleton Signaling | 9.64 |
| RhoGDI Signaling | 9.63 |
| G Beta Gamma Signaling | 9.38 |
| Role of Tissue Factor in Cancer | 9.33 |
| Ephrin Receptor Signaling | 8.93 |
| Signaling by Rho Family GTPases | 8.91 |
| Integrin Signaling | 8.34 |
| CXCR4 Signaling | 8.24 |
| Ephrin B Signaling | 7.93 |
| Virus Entry via Endocytic Pathways | 7.21 |
| Thrombin Signaling | 7.05 |
| Caveolar-mediated Endocytosis Signaling | 6.97 |

### Plasma HBPs

#### Identification of the plasma HBPs

Each plasma HBP isolation experiment was performed three times using blood obtained by cardiac puncture from control (NP) and AP mice. Acute pancreatitis was induced in animals as described previously. Histological and biochemical confirmation was consistent with the successful induction of AP (Supplementary figure 2).

The high abundance of serum albumin can pose problems for the analysis of plasma proteins by mass spectrometry. However, this protein does not bind heparin in the presence of electrolytes. To ensure initial binding of HBPs from plasma to heparin, the sample was applied to the heparin column in the lowest electrolyte concentration that prevented serum albumin binding. Thus, the citrated murine plasma was diluted with a PBS (150 mM NaCl) that was itself diluted with H_2_O to reduce the NaCl concentration to 75 mM. This was effective in reducing the albumin levels significantly in the eluate from the heparin column (Supplementary figure 3) and subsequent washing of the heparin column with PBS ensured that proteins bound heparin at physiologically relevant electrolyte concentrations. Thus, heparin affinity chromatography is a useful means to remove serum albumin from plasma samples, facilitating the MS analysis of plasma. Analysis of the MS data identified 161 plasma HBPs in NP (Supplementary Table 11) and 151 plasma HBPs in AP (Supplementary Table 12).

#### Label-free quantification

Label-free quantification was performed following the “Top3” methodology, as previously described (Fig. 6). This identified 69 HBPs that were overexpressed (Supplementary Table 13), which included known biomarkers of AP such as serum amyloid A (SAA) as well as emerging novel markers and drug targets in AP such as histone H2A (Table 5), and 81 that were underexpressed (supplementary Table 14) in AP. Interestingly, HRG, which was overexpressed in the pancreas in AP, was underexpressed in plasma in AP.

**Figure 6:**
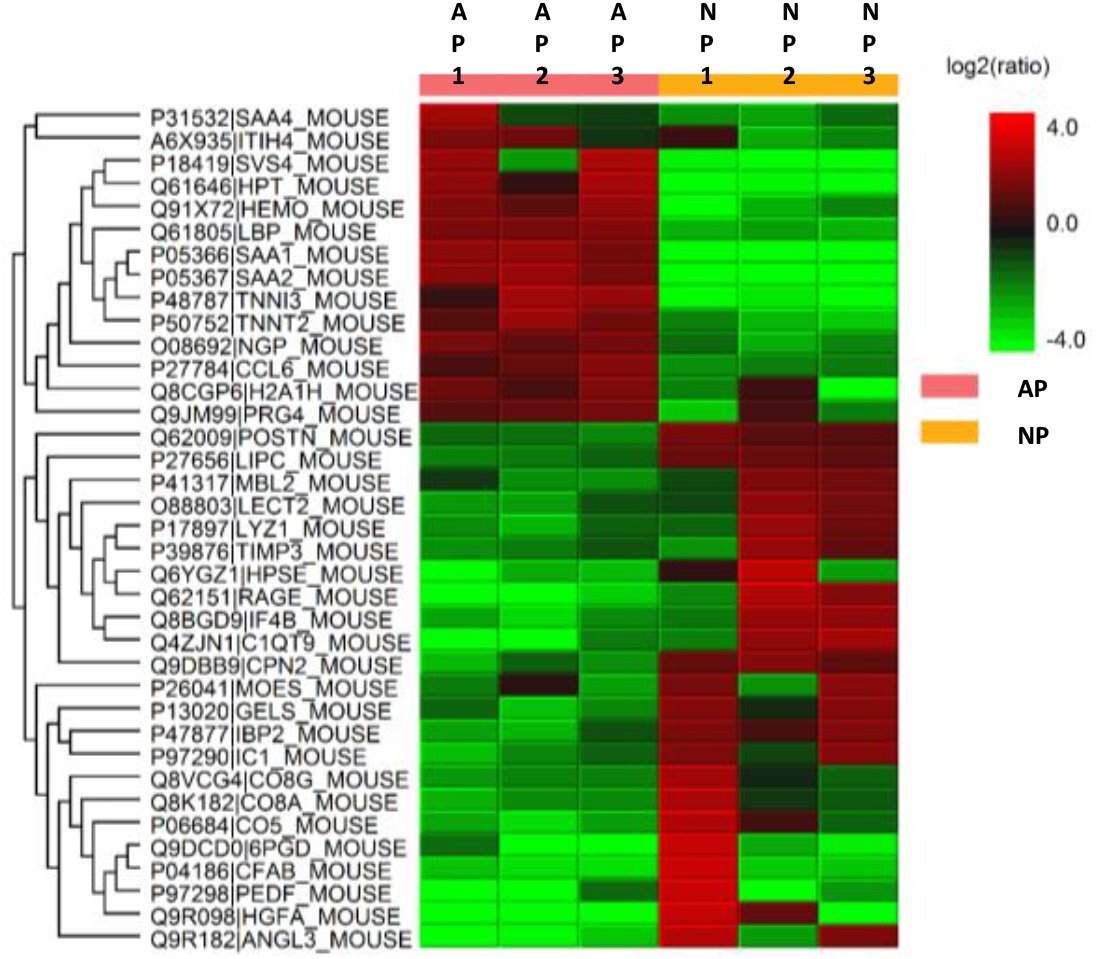
Heat map depicting the variation across the biological and technical replicates in plasma HBPs. The rows represent the various biological replicates from plasma in health (NP) and acute pancreatitis (AP), while the columns represent proteins. Red represents over expression and green represents under expression. Biological replicate number is denoted as “BioRep” and technical replicate number as “TechRep”. Hierarchical clustering was performed on both column data, to cluster the changes in protein expression, and row data, which displays the variation between samples.

**Table 5.** Top 20 Plasma HBPs overexpressed in AP. The upregulated HBPs were filtered depending on the maximum fold change values.

| **Plasma HBP** | **Max fold change** |
| --- | --- |
| SAA1 | 141 |
| Svs4 | 70 |
| HP | 15 |
| LBP | 6 |
| TNNI3 | 4 |
| Ngp | 3 |
| TNNT2 | 3 |
| ITIH4 | 2 |
| H2AFZ | 2 |
| Prg4 | 2 |

## Discussion

HBPs are functionally associated with the regulation of cell communication that underlies many physiological and pathological processes. AP represents a disease with unmet clinical need from the perspective of prognosis and treatment, particularly with respect to progression to severe AP, which can be life-threatening. Encouraged by the initial work using mRNA as a proxy for pancreas HBPs (13), we, therefore, determined whether there were changes in the levels of HBPs associated with AP in the murine caerulein model of the disease. One focus was on extracellular HBPs of the pancreas itself. Only extracellular HBPs were considered, because, whilst there is clear evidence for intracellular HS (42–44), there is no means at present to distinguish in a proteomic experiment intracellular proteins that interact with intracellular HS from those that natively interact solely with another intracellular polyanion, such as phosphorylated lipids and nucleic acids. The present analysis aimed to identify disease-related HBPs, and so processes, some of which may be potential therapeutic targets. However, potential biomarkers need to be more accessible, as sampling the pancreas is invasive, so plasma HBPs were also analysed. In this instance, because plasma is by definition cell-free, potentially intracellular proteins were also of interest. Moreover, the presence of exosome-associated proteins (supplementary Tables 3, 4, 11 & 12) is consistent with at least a subset of these vesicles binding heparin (45–47).

A number of tools were used to identify pancreas HBPs with significant amino acid sequence in the extracellular space, including SignalP, Phobius, SecretomeP, IPA, and manual curation for the presence of an extracellular signature using Uniprot-KB and PubMed. In a previous analysis of rat liver a bioinformatics pipeline based on ontology (gene ontology and ingenuity ontology, IPA) was combined with manual curation to identify 62 HBPs, of which 12 were previously known to bind to heparin (7). Middaugh and coworkers employing an antibody array on total cell lysate before and after depletion of HBPs by heparin agarose beads, identified 29 proteins whose signal was significantly reduced after incubation of the cell lysate with heparin beads, though only three were extracellular (48). Our present approach identified 360 new extracellular HBPs that were not previously known to bind the polysaccharide (7).

While plasma provides a rich and accessible repository of proteins, the large dynamic range of protein concentrations and high abundance proteins such as albumin pose a significant challenge to effective protein discovery (49, 50). However, albumin does not bind heparin in the presence of electrolytes and to ensure initial binding of HBPs from plasma to heparin, the sample was applied to the heparin column in the lowest electrolyte concentration that prevented serum albumin binding. In addition to extracellular/secreted proteins, a number of nuclear proteins such as histone H2A, cytoplasmic proteins such as FERMT3 (fermitin family member 3) and endoplasmic reticulum proteins such as HSPA5 (heat shock protein A5) were also detected in plasma.

An important question is how representative are the HBPs identified here of the proteins in NP and AP whose function depends on, or is modified by, interaction with HS. The mass spectrometry is limited by the depth of the analysis and only accepting identifications based on at least two peptides and a maximum false discovery rate of 1 %. This in itself leads to false positives and false negatives. To reduce further the number of false positives for the purpose of biomarker identification, stringency was increased by using the Bonferroni correction (21). There are additional factors that are likely to contribute to the false positives and false negatives. Some HBPs may escape detection, because they are part of large macromolecular assemblies that are not solubilised by Triton-X-100, associated with, for example, matrix fibrils, membrane microdomains or cytoskeleton. Moreover, not all HS-binding proteins will necessarily bind the heparin column and may instead remain bound to HS proteoglycans during affinity chromatography. Indeed, the transition and S-domains of HS have far more diverse structures than heparin (8, 9). Thus, the present analysis will have a bias towards HBPs that bind structures present in the trisulfated disaccharide repeat of heparin, which makes up 75% of the polysaccharide and contains 2-O sulfated iduronate, N-, 6-O sulfated glucosamine. A source of false positives would be proteins that do not bind HS, but are bound to an HBP with a sufficiently slow dissociation rate constant that they would be carried though the affinity chromatography, an interaction that would be consistent with biological function. Apart from the limits imposed on the mass spectrometry analysis, the other sources of false positives and negatives are not currently quantifiable, until such time as the interaction of each HBP with the polysaccharide is measured directly.

A recurrent question is how specific or selective is a protein-HS interaction (8, 9). This will be important in terms of understanding the mechanism whereby HS regulates the activity of a protein. HS has been shown to directly control the movement of a protein, fibroblast growth factor-2 in the pericellular matrix (51). Thus, regardless of specificity and selectivity of the interaction of a protein with HS, this will restrict the movement of the protein in the extracellular space and so affect its function in terms of location and local concentration. Therefore, proteins that bind heparin in PBS and that remain bound during the extensive washing of the affinity column have a relatively slow dissociation rate constant and would be expected to have at least their movement and extracellular location regulated by HS. Taking into account the caveats described above regarding false positives, the HBPs identified here are thus likely to have at least this aspect of their function regulated in this way by HS in the extracellular space of NP and AP. Any changes in the structure of HS that might accompany AP may, therefore, alter the movement of the HBP and so its contribution to cell physiology.

Thus, the heparin interactome is now substantially larger than previously described (7). Importantly, the protein-protein interaction network of the HBPs retains the key properties of the earlier, smaller interactome and hence its functional relevance to the regulation of cell communication by extracellular proteins (7). Thus, the HBP interactome is highly interconnected and HBPs form numerous regulatory modules in the extracellular space. Importantly, this is true for the subset of HBPs expressed by a single organ, the pancreas and one of its associated diseases, AP. Thus, it may be that for most if not all organs, HBPs represent a highly interconnected set of extracellular proteins, which have significant regulatory potential.

This work has identified a range of changes that provide deeper insight into known mechanisms in AP. A number of biomarkers that have been previously investigated in AP, such as carboxypeptidase B and pancreatic amylase, are amongst the HBPs that exhibited a significant increase in the extracellular pancreas AP dataset (Table 1 and Supplementary Table 5). Others such as SAA1 are significantly overexpressed in plasma in AP (52). Some of the most highly expressed pancreas HBPs such as NGP and HRG are representative of the underlying role of innate immunity in early AP. NGP overexpression indicates the early presence of neutrophils in AP (53), which have been shown to play an important role in AP (37). Neutrophil extracellular traps (NETs) are composed of extracellular DNA, neutrophil-derived granule proteins, and histones and are produced in AP as well as play a critical role in its development (54). HRG is an important mediator of macrophage clearance of apoptotic cells (38) and plays important roles in the adhesion and spread of activated T-cells (39). In contrast to its elevated levels in the pancreas, HRG is underexpressed in plasma in AP. Histone H2A levels were significantly increased in plasma in AP. The quantitative assessment of histones in plasma has been demonstrated to accurately predict persistent organ failure and mortality in AP (55). Thus, the HBPs exhibiting the greatest fold change (Table 1) alone or in combination with existing biomarkers may provide an improved clinical stratification of the severity of AP. Others such as serpin peptidase inhibitor, clade C (antithrombin), member 1 (SERPINC1) have anti-inflammatory properties and have been shown to improve acute pancreatitis in the rat model (56) and may be explored in drug development in AP. Some of the HBPs that are underexpressed in AP may point to a possible disturbance in the homeostatic mechanisms underlying AP. The underexpression of SLC4A7, KCNQ1 and VAMP2 (Table 2) are indicative of the impaired cellular trafficking of electrolytes that accompanies AP. In addition to its important role in K+ homeostasis, KCNQ1 also influences hormone transport (57).

CXCR4 signalling regulates cell differentiation, cell chemotaxis, cell survival and apoptosis, and has important roles in the embryonic development of the pancreas (58). Its importance as a key canonical pathway in AP is supported by the identification of its protective role in AP, which may be mediated by facilitating the migration of bone marrow derived stem cells towards the pancreas (59). Early activation of the ‘Complement system’, a pathway enriched to the extracellular pancreas HBPs in AP, occurs in pancreatic necrosis and corresponds to the overexpression of various proteins of the complement system in the plasma in AP, which suggests that this pathway may enable the development of treatment of leukocyte-associated injury in AP (60). In this respect, non-anticoagulant heparin-based compounds (61) might be particularly interesting as potential therapeutics for AP. They may be tuneable to modulate a gamut of HBPs important in the cell communication underlying the progression of AP, which could include complement system activation, growth factors and cytokines, and, because heparin can modulate the activity of some transporters (62), then metabolism.

The HBPs define a group of proteins that have clear functional importance in cell communication, homeostasis and at least one disease, AP. HBPs are experimentally accessible, because they bind to heparin and are extracellular, and in at least some instances are present in serum. Therefore the HBPs of the pancreas and AP may yield much-needed biomarkers and targets for therapy, a conclusion reinforced by the observation that HBPs identify through their functions to known mechanisms of AP and current targets for therapy in AP and other diseases.

## Ethics

The experiments were conducted in compliance with UK Home Office regulations (PL4003320), and were approved by the Institutional ethical review Committee of the University of Liverpool.

## Data accessibility

The mass spectrometry proteomics data have been deposited to the ProteomeXchange Consortium (63) via the PRIDE partner repository (http://www.ebi.ac.uk/pride/archive/) (64) with the following dataset identifiers:

1. Pancreas HBPs PXD001950
2. Plasma HBPs PXD012039

## Authors’ contributions

QMN isolated the plasma membrane fraction from pancreas, performed the functional analyses, drafted the paper; SD isolated the plasma HBPs, aided with the analyses and revised the paper; PJB and DS performed the mass spectrometry, aided with the analyses and revised the paper; CS performed the affinity chromatography experiments, aided with the analyses and revised the paper; YL performed the affinity chromatography experiments, aided with the analyses and revised the paper; TPB aided with analyses and revised the protocol and the paper, WH and XZ significantly contributed to the design of the animal experiments and revised the paper; DJR aided significantly with the bioinformatics analyses and revised the paper; RJB aided with the design of the mass spectrometry experiments and analyses and revised the paper; RS conceived the study and revised the paper; DGF conceived the study, co-wrote the paper and revised it All authors gave final approval for publication.

