## Supplemental Figures for "The heparin-binding proteome in normal pancreas and murine experimental acute pancreatitis"

### Slide 1
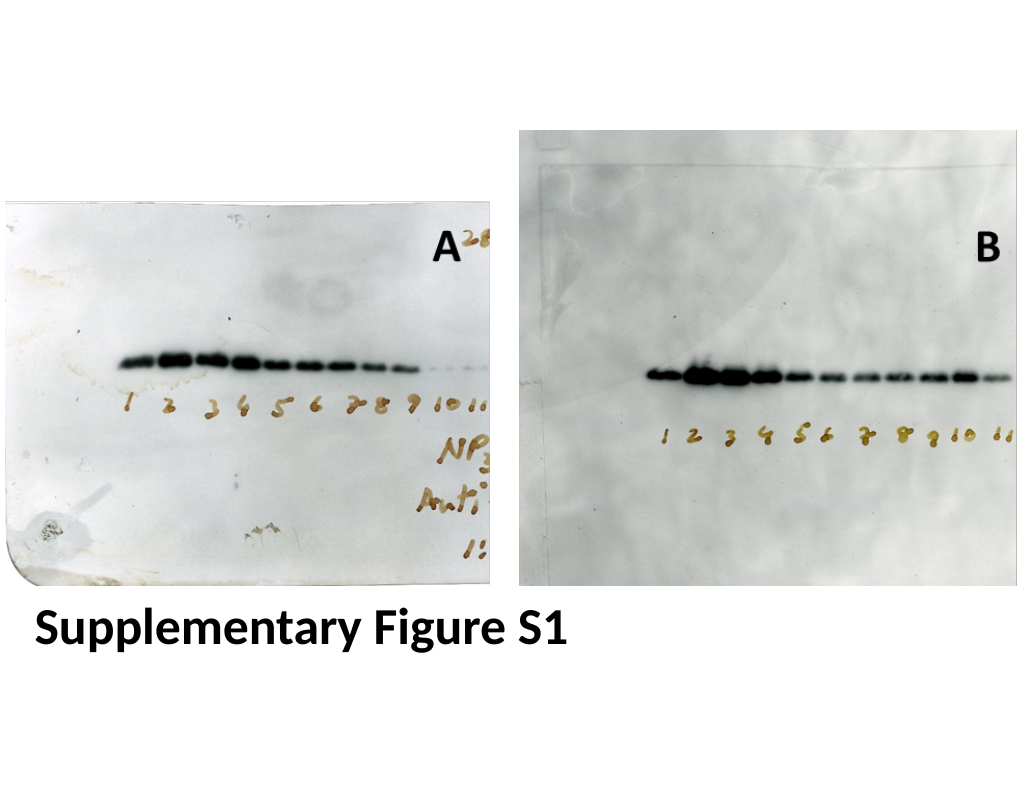

Supplementary Figure S1

### Slide 2
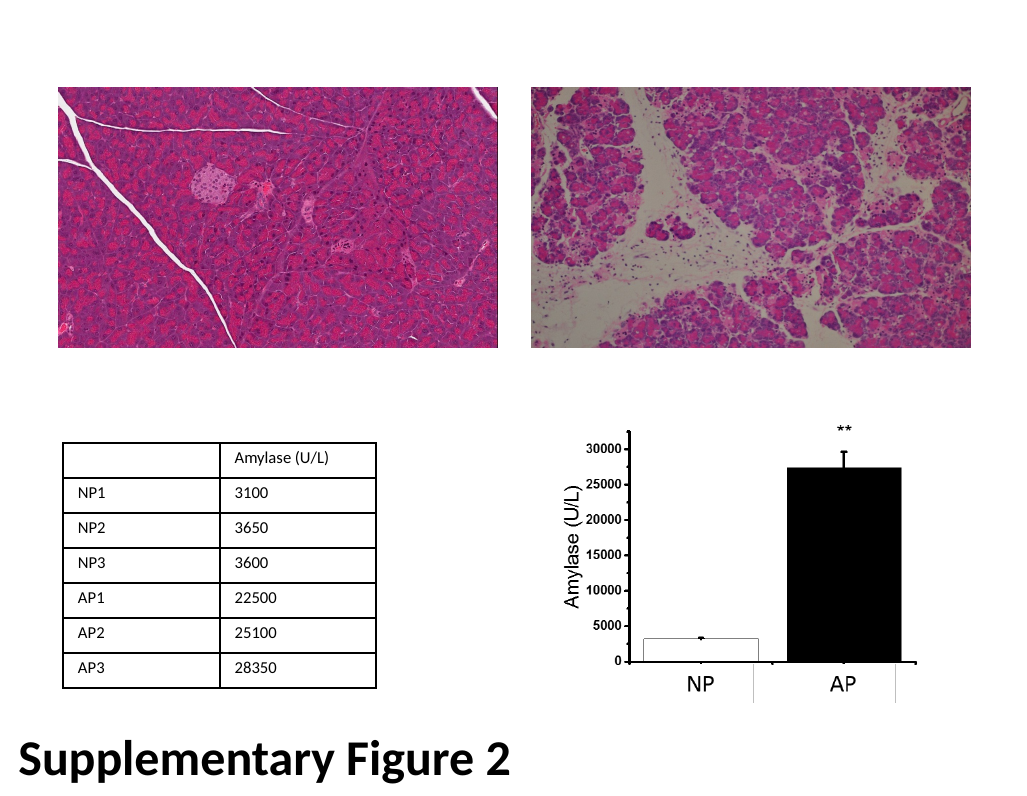

| | Amylase (U/L) |
| --- | --- |
| NP1 | 3100 |
| NP2 | 3650 |
| NP3 | 3600 |
| AP1 | 22500 |
| AP2 | 25100 |
| AP3 | 28350 |
Supplementary Figure 2

### Slide 3
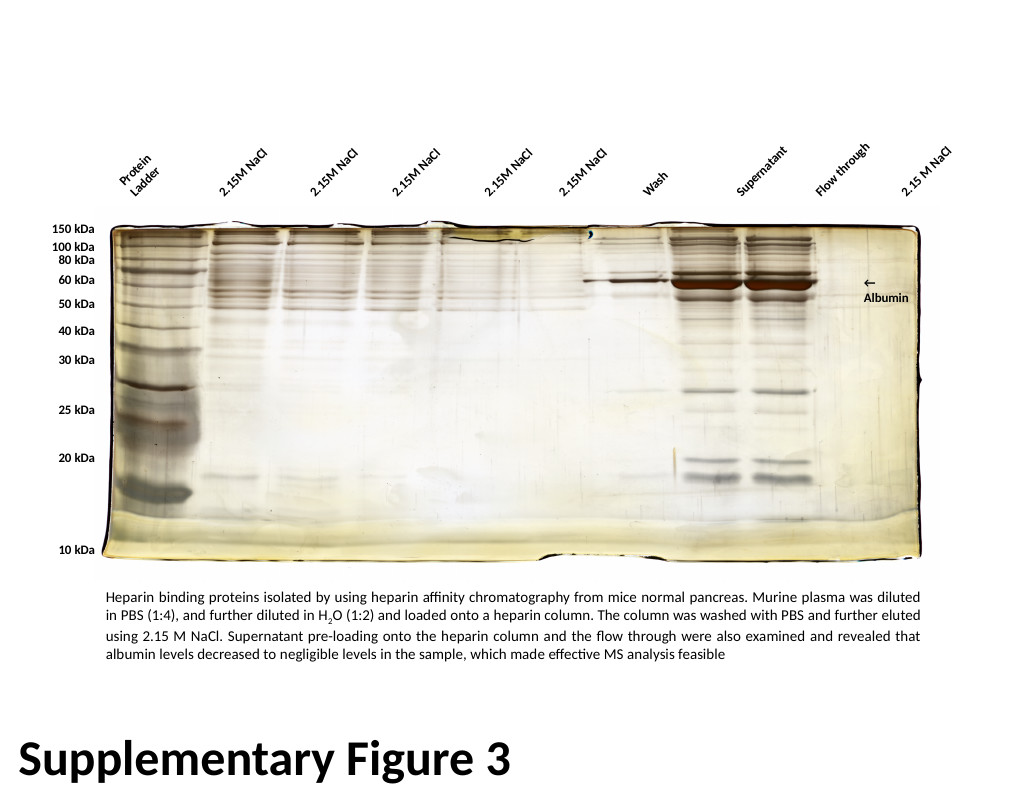

2.15 M NaCl
Supernatant
Protein Ladder
Flow through
2.15M NaCl
2.15M NaCl
2.15M NaCl
2.15M NaCl
2.15M NaCl
Wash
150 kDa
100 kDa
80 kDa
60 kDa
← Albumin
50 kDa
40 kDa
30 kDa
25 kDa
20 kDa
10 kDa
Heparin binding proteins isolated by using heparin affinity chromatography from mice normal pancreas. Murine plasma was diluted in PBS (1:4), and further diluted in H_2_O (1:2) and loaded onto a heparin column. The column was washed with PBS and further eluted using 2.15 M NaCl. Supernatant pre-loading onto the heparin column and the flow through were also examined and revealed that albumin levels decreased to negligible levels in the sample, which made effective MS analysis feasible
Supplementary Figure 3

### Slide 4
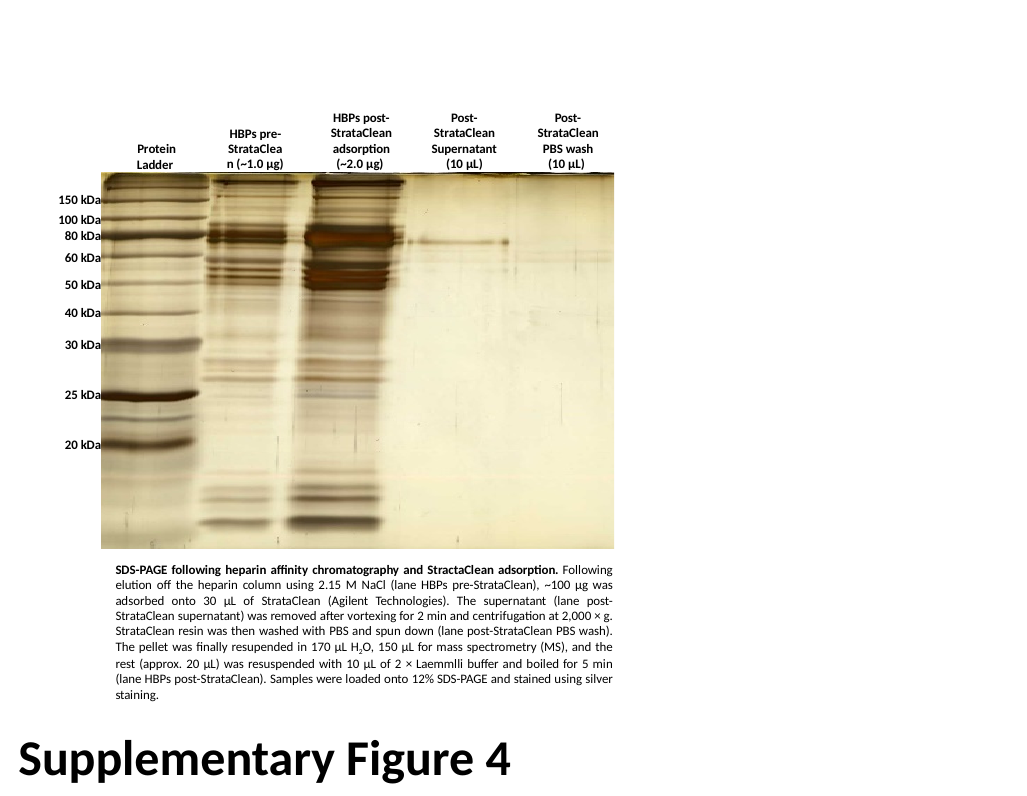

HBPs post-StrataClean
adsorption (~2.0 µg)
Post-StrataClean Supernatant (10 µL)
Post-StrataClean PBS wash (10 µL)
HBPs pre-StrataClean (~1.0 µg)
Protein Ladder
150 kDa
100 kDa
80 kDa
60 kDa
50 kDa
40 kDa
30 kDa
25 kDa
20 kDa
SDS-PAGE following heparin affinity chromatography and StractaClean adsorption. Following elution off the heparin column using 2.15 M NaCl (lane HBPs pre-StrataClean), ~100 µg was adsorbed onto 30 µL of StrataClean (Agilent Technologies). The supernatant (lane post-StrataClean supernatant) was removed after vortexing for 2 min and centrifugation at 2,000 × g. StrataClean resin was then washed with PBS and spun down (lane post-StrataClean PBS wash). The pellet was finally resupended in 170 µL H_2_O, 150 µL for mass spectrometry (MS), and the rest (approx. 20 µL) was resuspended with 10 µL of 2 × Laemmlli buffer and boiled for 5 min (lane HBPs post-StrataClean). Samples were loaded onto 12% SDS-PAGE and stained using silver staining.
Supplementary Figure 4
